# JADE1 and HBO1/KAT7 proteins in the cytokinesis of epithelial cells. The role of PHD zinc fingers

**DOI:** 10.1101/2022.10.13.512167

**Authors:** Bo Shao, Maria V. Panchenko

## Abstract

Members of the conserved subfamily, JADE1S and JADE1L isoforms, are expressed in epithelial cells, fibroblasts, and epithelial cell lining in vivo. JADE1 proteins interact with histone acetyl transferase HBO1 complex. The two consecutive PHD zinc fingers of JADE1 bind chromatin. We recently reported novel effects of JADE1S on cytokinesis progression. JADE1S depletion facilitated G2/M-to-G1 transition and increased polyploidy and aneuploidy. JADE1S over-expression arrested cells in late cytokinesis, an effect reversed by AURKB inhibitor. In late cytokinesis cells JADE1S protein localized to the midbody. Results suggested a JADE1S role in final abscission delay. Here we investigated the expression of JADE1 in the central spindle, interactions with HBO1, and the role of PHD fingers in late cytokinesis arrest.

The midzone begins to assemble in anaphase and forms into a midbody in cytokinesis. The midbody structure connects two daughter cells and is thought to bear factors controlling the final abscission. We questioned whether, similar to established factors, JADE1S is targeted to the central spindle structures in anaphase. Indeed, in cells transitioning from mitosis to cytokinesis, JADE1S was sequentially targeted to early midzone, midbody flanking zone, and midbody. The step-wise increase of JADE1S expression in midzone and midbody of synchronously dividing cells suggested protein recruitment. The increase of late cytokinesis arrest caused by recombinant JADE1S correlated with increased expression in midbody. Spatial analysis of the members of the chromatin passenger complex, microtubule associated proteins, and centralspindlin, revealed transient co-localization with JADE1S and mapped JADE1S within the cytokinesis bridge.

Deletion of the two PHD zinc fingers inactivated JADE1S ability to arrest cells in late cytokinesis but did not affect its midbody localization. Thus, PHD zinc fingers are required for JADE1S cytokinesis delay but not for midbody targeting. Recombinant HBO1 protein decreased the proportion of late cytokinesis cells, prevented late cytokinesis arrest by JADE1S as well as its midbody localization. Enzyme inactive HBO1 mutant recapitulated the wild type phenotype. The results demonstrate antagonistic relationship and suggest HBO1-mediated midbody dislocation of JADE1S. Our study supports the role of JADE1S in cytokinesis delay and implicates protein partners.

## Introduction

The Gene for Apoptosis and Differentiation in Epithelia-1 (JADE1/PHF17) belongs to the small family of JADE proteins and is highly expressed in a variety of cultured epithelial cells as well as in epithelial cell lining of several organs in mammals[1-9].

The JADE1 gene gives rise to two protein products, JADE1S and JADE1L isoforms. The C-terminal of JADE1S protein is missing the 333-amino acid fragment of the full length isoform and includes a unique stretch of seven C-terminal amino acids. The remaining sequence of 502 amino acids in JADE1S polypeptide is identical to JADE1L. Both JADE1 isoforms bear one canonical Cys4HisCys3 plant homeodomain (PHD) followed by a non-canonical extended PHD domain, which are zinc-binding motifs[10]. We and others previously reported results suggesting functional differences between the two JADE1 isoforms[4-7, 11] and (unpublished). The cellular function of JADE1L has not been elucidated.

Recent studies implicate the JADE1S role in the cell cycle[4, 5, 7, 9, 12]. The JADE1S protein is dynamic and undergoes chromatin-cytoplasm shuttling during mitosis which is linked to its phosphorylation[9]. Six phosphorylated amino acid residues in mitotic specie of JADE1S were identified by Mass Spectroscopy analysis[9]. The chromatin dissociation and phosphorylation of JADE1S were prevented by the pharmacological inhibitor of Aurora A kinase[9]. Depletion of JADE1S protein in cell cycle synchronized cell cultures increased the rates of G2/M to G1 transition[5]. In randomly dividing cells, JADE1S depletion increased polynucleation and aneuploidy while JADE1S overexpression increased proportion of cells in late cytokinesis. The apparent arrest of cells in late cytokinesis mediated by JADE1S over-expression was released by the addition of AURKB inhibitor[5]. Based on these and other results we suggested JADE1S function in AURKB-mediated final abscission delay.

Previous studies characterized activities of JADE1S in the nucleus [3, 4, 7, 9, 12-17]. Thus, JADE1 protein is associated with chromatin and is a candidate transcription factor[3]. JADE1 promotes histone H4 acetylation by associating with a histone H4-specific endogenous HAT in cultured cells and in vitro[3]. In the context of chromatin, the histone acetylation activity of JADE1 requires intact PHD zinc fingers, suggesting a chromatin-targeting role for PHD zinc fingers in live cells[3, 7, 11, 16]. JADE1 has been characterized as a component of the HAT HBO1 complex which is the most studied protein partner[3, 4, 7, 9, 11, 14, 16, 18]. Other protein partners of JADE1 have been also reported[2, 12, 15, 17, 19].

HBO1 (MYST2, KAT716) was originally identified in a yeast two-hybrid screen as a HAT binding origin recognition complex-1 (Orc1)[20, 21]. Studies implicate the HBO1 role in pre-replication complex assembly, DNA synthesis, transcriptional regulation as well as cellular stress response and carcinogenesis[11, 16, 22-29]. A recent study reported that HBO1 HAT activity positively affected centromeric CENP-A assembly, presumably via increased histone turnover and interactions with other proteins regulators of centromere assembly and function[30]. The authors suggested a contribution by another partner of HBO1, PHD zinc finger protein and JADE1 homolog, BRPF1. The cooperative interactions of JADE1 with the components of the HBO1 complex have been established[4, 7, 9]. In addition to JADE1, the cellular activities of the HBO1 complex might be controlled by the presence of other PHD zinc finger bearing partners[11, 14, 18, 31].

Cells enter cytokinesis following chromosome segregation in anaphase. The beginning of cytokinesis is manifested by cleavage furrow ingression which is driven by actomyosin contractile ring. The dynamic microtubule network is essential for cytokinesis progression and control. An important microtubule-based structure, central spindle, assembles between segregating chromosomes in anaphase. The plus-ends of interpolar antiparallel microtubule fibers overlap in the middle of the central spindle, or the midzone. Central spindle and astral microtubules network define the cleavage furrow central positioning. As cytokinesis progresses the constricting actomyosin ring tightens around the central spindle bundle which eventually matures into a structure called midbody. The assembly of midbody has been under investigations. A large number of specialized proteins are recruited to midbody from various cellular compartments prior to the final abscission and contribute to cytokinesis control and progression. The endosomal sorting complex required for transport (ESCRT) machinery contributes to midbody assembly and final abscission by recruiting other factors and forming scission filaments[32-35]. Chromatin passenger complex (CPC) proteins, microtubule-associated proteins (MAPs), and centralspindlin complex are targeted to the central spindle to contribute to its assembly and stabilization, as well as furrow ingression timing in early cytokinesis[33, 36-43]. In late cytokinesis, as cells prepare for final abscission, several of these factors consolidate in the flanking midbody zone as well as inside the midbody (Flemming body) [38]. It has been proposed that midbody serves as a platform for control and execution of final abscission. At least two known complications of mitosis, such as the presence of lagging chromatin fibers trapped within the cytokinesis bridge and defects with nuclear pore reassembly, may result in the activation of the final abscission delay, also called cytokinesis checkpoint, NoCut [34, 44-47]. Failure to activate the cytokinesis checkpoint may result in polyploidy, aneuploidy, apoptosis, as well as DNA loss and damage, all of which have been considered as hallmarks of cancers. Factors that activate final abscission delay are under active investigations.

In the current study we further investigated the role and regulation of JADE1 during cytokinesis progression. We performed spatial-temporal analysis of JADE1 expression starting from mitotic exit and lasting to late cytokinesis. We mapped JADE1S localization within the central spindle structures relative to established midzone/midbody proteins and cytokinesis regulators. Our results show that targeting of endogenous JADE1S to midbody occurs via central spindle midzone and implicate stepwise recruitment of JADE1S. We identified the requirement of JADE1 PHD zinc finger module in recombinant JASE1S-mediated cytokinesis arrest, and determined the competitive HBO1 effects on JADE1S-mediated late cytokinesis arrest and midbody targeting. Our study signifies the role of JADE1S in late cytokinesis arrest and suggests interactions with established factors of cytokinesis.

## Materials and methods

### 1. Cell culture

HeLa, H1299, and HEK-293 were obtained from ATCC. All cells were grown in Dulbecco’s modified Eagle’s medium supplemented with 10% (v/v) fetal bovine serum and 1% (v/v) Penicillin-Streptomycin (Cellgro), at 37^0^C in a humidified incubator with 5% CO2 atmosphere. Sub-confluent cells grown in 60-, 100-mm dishes, or 2-well and 4-well chamber slides were transiently transfected with various cDNA constructs using Lipofectamine 2000 (Invitrogen) or FuGENE 6 (Promega) following the manufacturer’s protocol with modifications (see below).

### 2. Pharmacological treatments

Cells were treated with drugs and cell cycle profiles characterized according to the previously published protocol[5, 9]. Briefly, to synchronize cells in G1/S phase, L-mimosine (250 μM) was added to cell cultures for 16 hours. Cell cycle was released by replacing the conditioning media with fresh media. After release into fresh media, the cells were processed for immunofluorescence at different time points as indicated.

### 3. Protein extraction from cultured cells

Cells were collected in cold PBS, and centrifuged at 1000 rpm for 5 min at 4°C (accuSpin Micro 17R; Fisher Scientific). The proteins were extracted in 50 mM Tris buffer (pH 7.8) containing 5 mM MgCl2, 0.5% NP-40, 150 mM NaCl, 1 mM EDTA, protease and phosphatase inhibitors and PMSF (10 minutes on ice) and centrifuged at 13,300 rpm for 5 min at 4OC (Centrifuge 5810 R; rotor A-4-81; Eppendorf). The supernatant (whole cell lysates) was analyzed using western blot analysis.

### 4. Antibodies and chemicals

Mouse monoclonal antibodies for FLAG M5 (1:1000; F1804), alpha-tubulin (1:1000; T9026) and beta-actin (1:250; A1978), as well as rabbit polyclonal antibodies for FLAG (1:1000; F7425), were from Sigma-Aldrich. Mouse monoclonal antibodies for alpha-tubulin (1:1000, ab18251), HBO1 (1:250; ab70183), and AURKB (1:700, ab3609) were from Abcam. Mouse monoclonal antibody for gamma-tubulin (1:300; MA1-850) was from Thermo Scientific. Mouse monoclonal antibodies against PRC1 (1:1000, sc-376983), KIF4A (1:500, sc-365144), MgcRacGAP (1:500, sc-271110), and Goat anti-mouse and anti-rabbit IgG-horseradish peroxidase conjugates (1:2000; sc-2005 and sc-2004) were from Santa Cruz Biotechnology. Alexa Fluor dye-labeled secondary antibodies (1:500; A31627, A31623, A31619, A31631) were from Life Technologies. Protease inhibitor cocktail and phosphatase inhibitor cocktail was from Roche Diagnostics. L-Mimosine (M0253) was from Sigma-Aldrich.

### 5. Custom made antibody specificities

The specificities of the custom made polyclonal sera and affinity-purified HBO1-, JADE1L-, and JADE1S-antibody reagents have been examined in various assays, validated, and documented in our recent publications [5, 9]. To achieve specificity and avoid cross-reactivity between JADE1S and JADE1L isoforms we designed two unique peptides as antigens for either antibody production: JADE1S antigen included unique C-terminal peptide (RQDLERVMIDTDTL), while JADE1L antigen peptide was chosen from the unique extra C-terminal of the protein (LKSDNENDGYVPDVEMSDSE). Sera were further purified by affinity chromatography against the corresponding peptides by the vendor. Experimental validation and specificity controls for antibodies have been recently described in detail by using protein depletion as well as protein detections in various cell fractions[5, 9] (Siriwardana et al, 2015, as examples see Figure 1, D; Figure 8 A and B). Additional validation of JADE1 antibodies in the current study shows no protein isoform cross-reactivity (this study, Results section).

**Fig 1.**
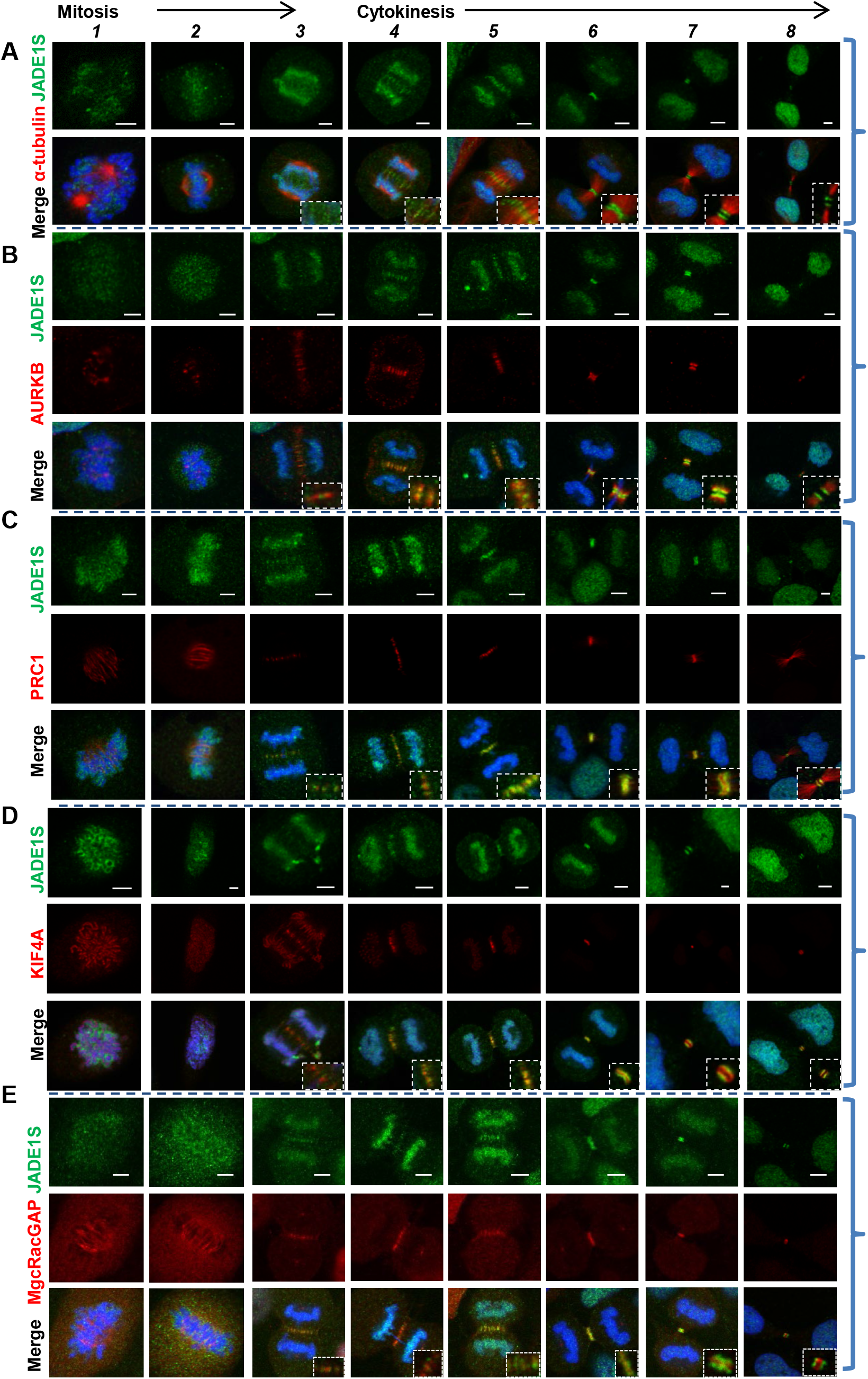
JADE1S transitions from early midzone to late midbody and partially co-localizes with key factors of mitosis and cytokinesis. **(A-E)** Asynchronously dividing HeLa cells were processed for immunofluorescence (IF) and proteins visualized with antibodies as indicated. DNA is visualized with DAPI. Representative confocal images of cells transitioning from mitosis to late cytokinesis. **(A)** Endogenous JADE1S is concentrated to early forming midzone in late anaphase (#*4*), to midzone in early cytokinesis and midbody in late cytokinesis (#*5* to #*8*). **(B-E)** Spatial-temporal coordination of JADE1S expression with (B) AURKB, (C) PRC1, (D) KIF4, and (E) MgcRacGAP. Scale bar, 5 μm

### 6. Constructs of cDNA used in the study

HA- and FLAG-tagged JADE1S, HA-JADE1dd, HA- and FLAG-tagged wild type and E/Q mutant HBO1 were described previously [3, 5, 7, 9]. FLAG-JADE1L was described in[5]. MGC (Mammalian Genome Collection) fully sequenced human PHF17 (JADE1L) cDNA cloned in pCMV-SPORT6 vector (clone 5111727) was obtained from Open Biosystems. PHF17 cDNA was then cloned into pcDNA3.1 vector modified with the N-terminal FLAG-tag.

### 7. Transfection and immunofluorescent labeling of whole cells

Immunofluorescence (IF) was done as described, except for the following modification[5, 9]. Cells grown on two-or four-well chamber slides for 36-48 hours were transfected with Lipofectamin 2000 or FuGENE. To ensure low physiological levels of protein expression and avoid toxicities, the quantities of cDNA and transfection reagents were scaled down and did not exceed one half or, in some cases, one third of that recommended by manufacturers. Cells were allowed to grow for 24-26 hours and processed for immunofluorescence by rinsing with PBS, and then fixing with 4% paraformaldehyde for 15 min at room temperature. Cells were rinsed with PBS three times and then permeabilized with 0.5% or 3% Triton X-100 for 15 min at room temperature. Blocking was performed in 0.05% Tween-PBS with 2% BSA for 2 hour at room temperature. Primary antibodies in 2% BSA, 0.05% Tween-PBS, were incubated overnight at 4°C and secondary antibodies in 2% BSA, 0.05% Tween-PBS were incubated for 1 hour at room temperature. Coverslips were mounted using Vectashield (VECTOR) mounting medium with DAPI. Images were analyzed on Leica TCS SP5 confocal system with an acousto-optic beam splitter for rapid switching between lasers and conventional point-scan as well as high-speed resonance scanning (Leica Microsystems, Cellular Imaging Core in Boston University School of Medicine) using 60X magnification lens. Images were edited in ImageJ 1.45S (NIH, Bethesda, MD) software. All experiments were repeated at least three times and in two different types of cells specified, yielding essentially the same results.

### 8. Quantitation of mitotic, cytokinetic, and multinucleated cells. Plot profile analysis of cytokinesis bridge

Cells were scored by manual counting under the fluorescence microscope (Olympus BX60) to quantify the cytokinetic, bi- and multinucleated cells as percentage of total transfected cells. When specified, mitotic and cytokinesis cells were quantified and analyzed within each sub-population, with representative images provided. When indicated the total number of cells analyzed is provided in Legends to Figures. All experiments were repeated three times and data are presented as average ± SD. To calculate the fold change the percentage of cytokinetic cells at each data point values were normalized to the value of the first data point. Plot profile analysis of the central spindle was done using ImageJ 1.45S (NIH, Bethesda, MD) software.

## Results

### 1. JADE1S is targeted to the early midzone in anaphase

According to our recent study, JADE1S over-expression resulted in increased proportion of late cytokinesis cells, indicating cytokinesis arrest, while JADE1 depletion -in cell polyploidy. Depletion of JADE1S protein in cell cycle synchronized cell cultures increased the rates of G2/M to G1 transition suggesting role in late cytokinesis control[5]. Moreover, in late cytokinesis cells the nuclear chromatin associated protein JADE1S was localized to the midbody[5]. These results raised a possibility that, during cytokinesis progression, similar to other late mitosis and cytokinesis factors, JADE1S might be recruited to the midbody via the central spindle. Thus, we performed comprehensive analysis of cells exiting mitosis to determine if JADE1S is found within the forming central spindle microtubule structures. In early anaphase, which times to the beginning of the spindle growth and stabilization, a diffused amorphous density of JADE1S protein was detectable between the separating sister chromatids(Fig 1, A, *3*). At this point JADE1S was not visibly associated with microtubule fibers. In later anaphase cells, JADE1S concentrated to the middle of the central spindle, early midzone (Fig 1, A, *4*). This localization became more prominent upon furrow ingression in early telophase cells at which point JADE1S localized to the area of interdigitated tubulin plus ends (Fig1, A, *5*). In late telophase and consequently in cells with delayed cytokinesis (judged by elongated cytokinesis bridges) JADE1S concentrated to highly condensed tubulin fibers emanating from the midbody (Fig 1, A, *6-8*). JADE1S staining at the midbody flanking zone was bright and easily detectable. We sparingly detected the endogenous JADE1S protein inside the midbody. This is likely due to the antigen masking within the highly dense, tightly packed structure in midbody center (Flemming body, discussed further in Figs 4 and 5). Thus, similar to several midbody proteins[36], prior to its midbody localization in late cytokinesis, JADE1S associates with the antiparallel microtubule overlaps of the growing central spindle, or the early midzone, suggesting active recruitment.

### 2. Transient co-localization of endogenous JADE1S and CPC protein, AURKB

The cytokinesis arrest mediated by recombinant JADE1S was released with the addition of a pharmacological inhibitor of AURKB, suggesting a role in final abscission delay[5]. AURKB is one of the key kinase required for cytokinesis progression, coordination, and cytokinesis checkpoint, and is a catalytic component of the chromosome passenger complex (CPC) [42, 47-50]. The well-studied spatial temporal regulation of AURKB in mitosis includes the initial concentration of this kinase on centromeres in metaphase, followed by the recruitment to the central spindle midzone in anaphase and, finally, recruitment to the midbody flanking zone in late telophase and late cytokinesis[42]. We investigated whether JADE1S was colocalized with AURKB at any point in cells exiting mitosis and entering cytokinesis (Fig 1, B). As expected in metaphase cells AURKB localized to the characteristic speckles along the metaphase plate, marking centromeres prior to anaphase engagement (Fig 1, B, *1* and *2*). At this stage we did not detect any spatial associations between AURKB and JADE1S. With anaphase engagement and central spindle growth, AURKB concentrated to the assembling midzone, marking interdigitated tubulin fiber bundles between the two dividing cells (Fig 1, B, *3*). Already at this stage JADE1S protein concentrated and positioned immediately distal to AURKB. Strikingly, in early telophase (judging by constricting furrow and still condensed chromatin) JADE1S and AURKB became colocalized (Fig 1, B, *4*). JADE1S and AURKB partial colocalization was also evident in late telophase cells (Fig 1, B, *5-7*). Finally, in late cytokinesis cells with apparently delayed abscission (judging by the decondensing chromatin and the long bridge) JADE1S returned to localization immediate distal to AURKB (Fig 1, B, *8*). Our results demonstrate transient co-localization of JADE1S and AURKB in cells undergoing cytokinesis, suggesting interactions and further supporting the role in final abscission delay.

**Fig 2.**
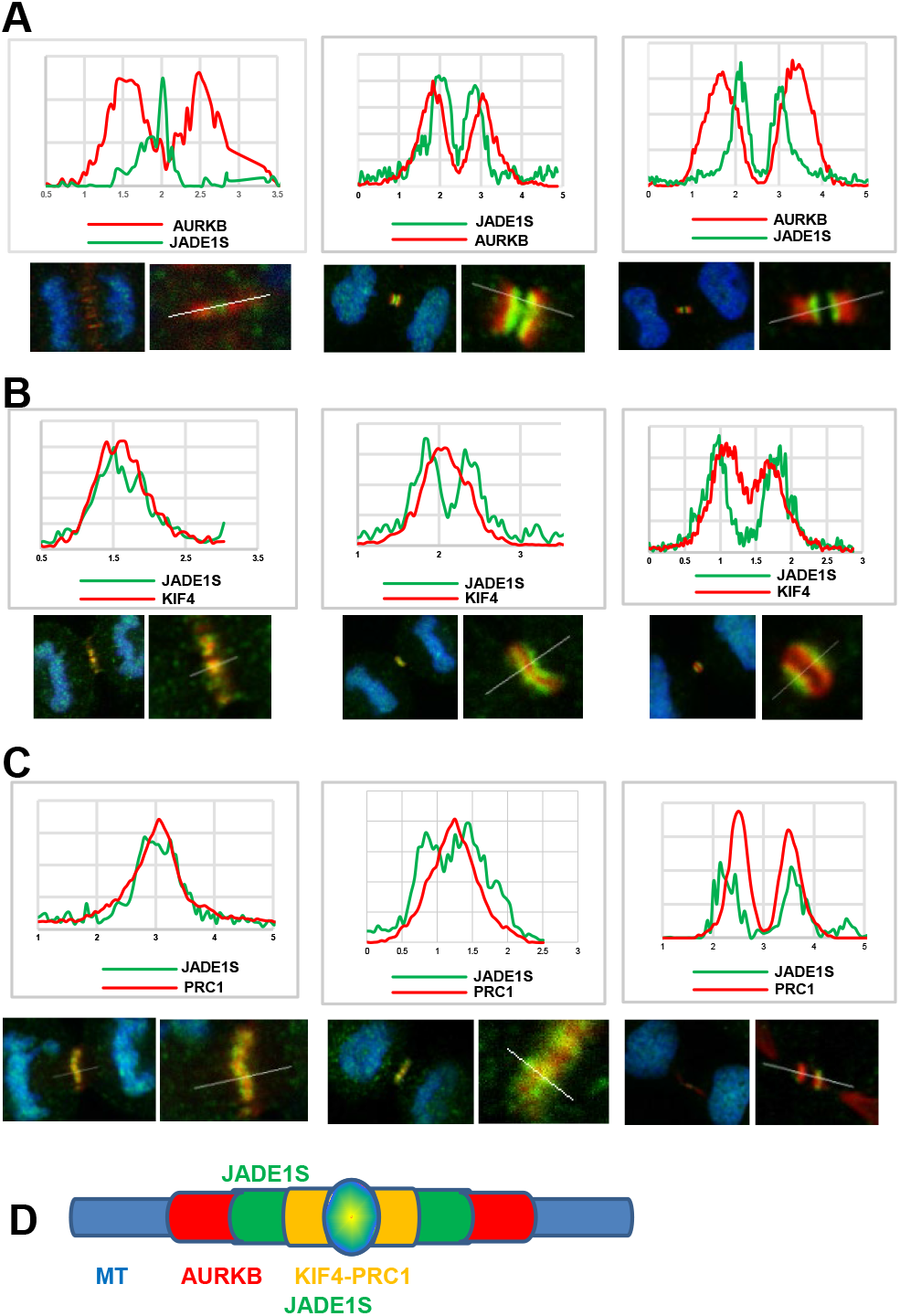
Plot profile JADE1S relative to CPC and MAP components in cells transitioning from mitosis to cytokinesis. **(A-C)** Plot profile analyses of central spindle in cytokinesis define midzone and midbody JADE1S position relative to CPC and MAP components (A) AURKB, (B) KIF4A, and (C) PRC1. **(D)** Schematic summary of (A-D) illustrates JADE1S relative positioning in late cytokinesis central spindle. Results supporting JADE1S co-localization with KIF4-PRC1 are presented in Fig 4.

**Fig 3.**
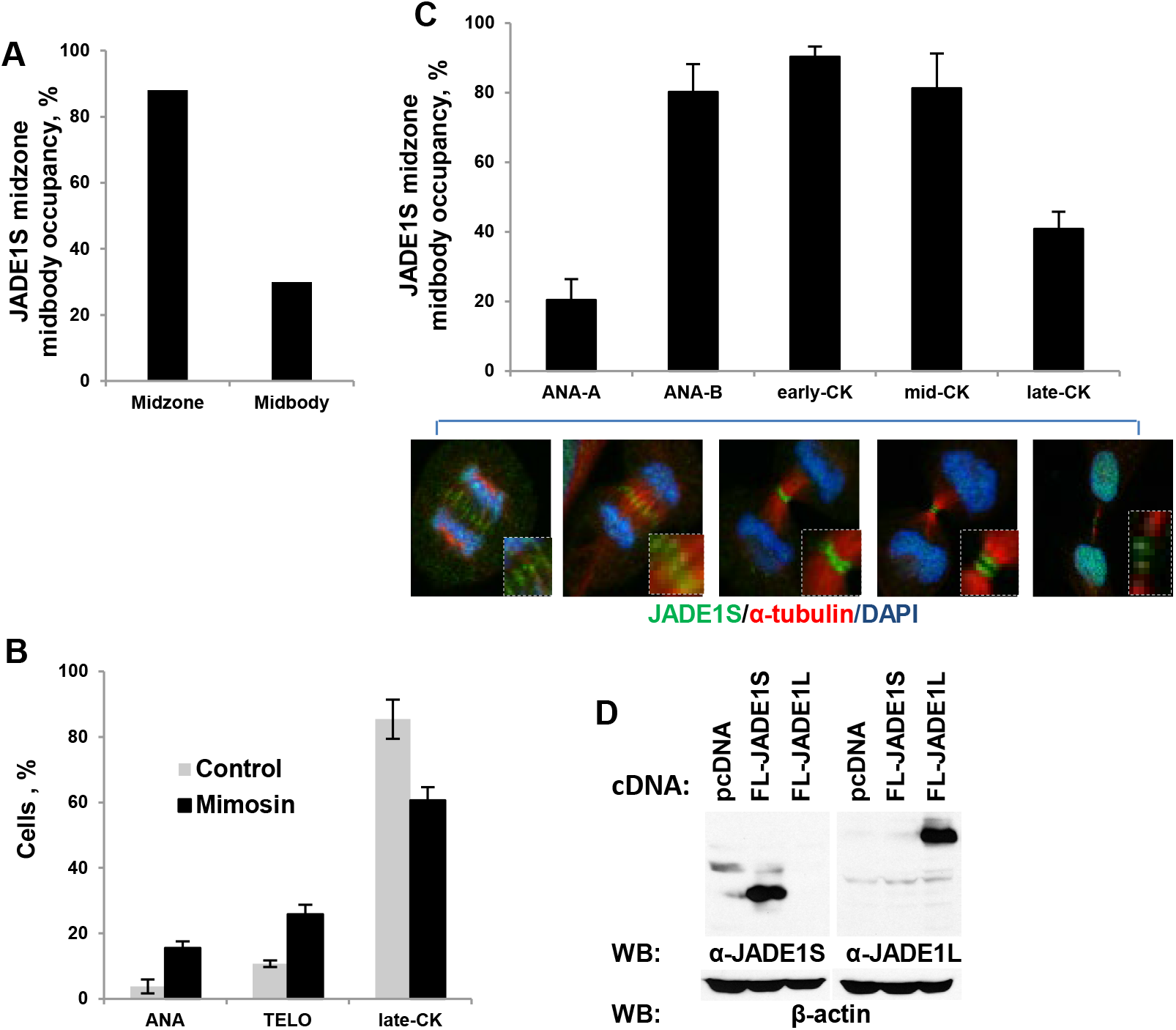
Endogenous JADE1S is recruited to the midzone and midbody in synchronized dividing cells. **(A) JADE1S midzone and midbody occupancy in asynchronous cells.** JADE1S and α-tubulin visualized by IF. All anaphase and cytokinesis cells were counted and divided to subpopulations according to their morphology as in (C): anaphase A (ANA-A), anaphase B (ANA-B), early cytokinesis (early-CK), mid-cytokinesis (mid-CK), and late cytokinesis (late-CK). To determine percent of JADE1S midzone occupancy (A, left bar), total population of cells with midzone in central spindle (ANA-B, early-CK, and mid-CK cells) was counted and percentage of cells positive for midzone JADE1S calculated. Total of at least 200 cells were analyzed per experiment. Similarly, to determine JADE1S midody occupancy (A, right bar), percentage of cells positive for midbody JADE1S was calculated in all late-CK population (>150 late-CK cells, reproduced twice). **(B) Cell cycle profile of cells transitioning from mitosis to cytokinesis after cell cycle release**. After 18, 20, and 22 hours of mimosine withdrawal and cell cycle release (details in[5,9]), proteins visualized by IF. Population of anaphase and cytokinesis cells was quantified and plotted as fraction of a total of ANA and CK combined. **(C) JADE1S midzone and midbody occupancy increases from ANA-B through mid-CK**. Cells were treated with mimosine and cell cycle released as in (B). Anaphase and cytokinesis cells were manually assessed according to their morphology as described in (A), representative images (C). Fraction of cells positive for JADE1S in the midzone, midbody flanking zone, or midbody were quantified within each of the sub-population and plotted. **(D) JADE1S antibody does not recognize JADE1L protein, and vice versa**. Western blot detection of recombinant proteins as labeled with antibodies specific for JADE1S (left panel) or JADE1L (right panel).

**Fig 4.**
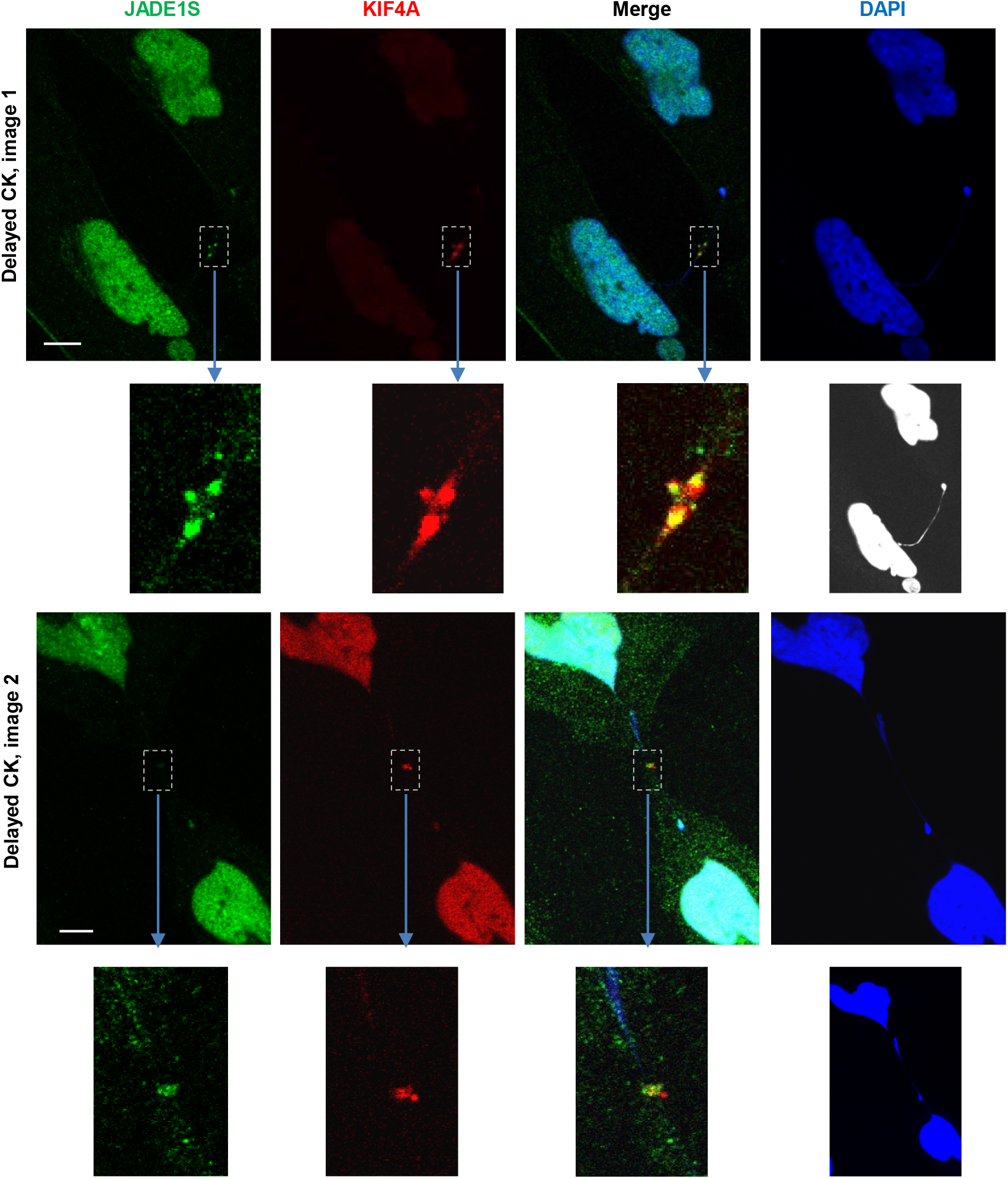
JADE1S and KIF4A are co-localized inside of the midbody (Femming body). Note that Flemming body JADE1S-positive cells have distinctive chromatin bridges detected by DAPI (discussed in Results and Discussion). Image #1 DAPI is in white to show the bridge, the inserts are adjusted bright to show detail. The composite for Image #2 is adjusted brighter green to show the Flemming body JADE1S detail. Bar 5μM

**Fig 5.**
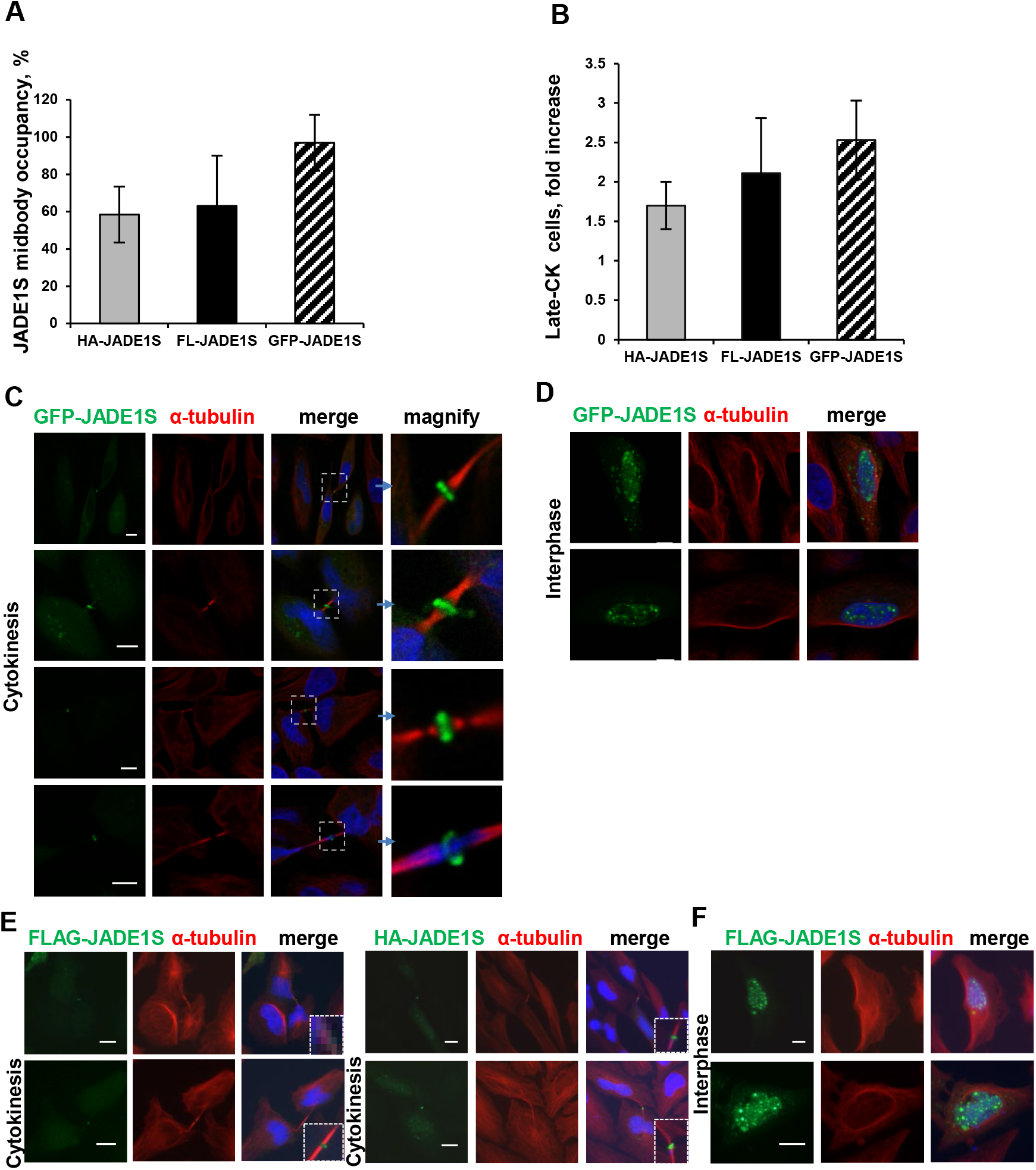
GFP-JADE1S is targeted to inside of midbody in majority of late cytokinesis cells. **(A) GFP-JADE1S midbody occupancy.** HeLa cells grown in 4-well chamber slides were transfected with 0.4 μg of either of the JADE1 cDNA: FL-JADE1S, HA-JADE1S, or GFP-JADE1S. Proteins visualized by IF with FLAG-or HA-monoclonal antibody in combination with rabbit α-tubulin antibody. In each well the total number of late cytokinesis cells expressing recombinant protein was counted, proportion of cells positive for midbody JADE1S quantified (midbody occupancy). Values are given as a percentage of the total transfected late cytokinesis cells from four independent experiments ± s.d., >200 cells counted per experiment.. **(B) GFP-JADE1S is the most potent in arresting late cytokinesis**. Experimental format, conditions, and quantitation as in (A), except empty vector control sample was included. Plasmids encoding HA-JADE1S, FLAG-JADE1S, GFP-JADE1S, or empty vector control were transfected to the cells and JADE1S-positive late cytokinesis cells were quantified as in[9]. Values represent fold increase of late cytokinesis cells over empty vector control (cells transfected with empty vector produced total of 10%±2 of late cytokinesis cells). (A,B) Error bars represents standard deviation. Note the correlation between tagged-JADE1S midbody occupancy (A) and cytokinesis delay effect (B). **(C) GFP-JADE1S concentrates exclusively inside the midbody in late cytokinesis cells**. Conditions as in (A) except half of the cDNA quantities were used. Note that GFP-JADE1S is found in the midbody with and without unresolved chromatin bridge. (**D**) **GFP-JADE1S in interphase cells is localized predominantly to the nuclei**. (**E**) **FLAG- and HA-JADE1S in the late cytokinesis midbody and (F) in interphase nuclei**. Representative images. Scale bar,10 μm.

### 3. Spatial-temporal expression of endogenous JADE1S, MAPs, and centralspindlin

In addition to the major CPC component, AURKB, we mapped the spatial-temporal expression of endogenous JADE1S relative to three specific factors of the central spindle assembly: (1) a highly conserved MAP called Protein Required for Cytokinesis-1 (PRC1)[51, 52]; (2) a direct partner of PRC1, kinesin KIF4[53, 54]; (3) centralspindlin component, MgcRacGap1[36, 55]. PRC1 binds microtubules, localizes to the central spindle and, in vitro, bundles microtubules, while KIF4 is required for PRC1 dimerization. Centralspindlin is a motor tetrameric complex made out of a dimer of the kinesin 6 motor protein MKLP1 and a dimer of the Rho family GTPase-activating protein MgcRacGap1[56]. This complex localizes to the middle of the central spindle, promotes central spindle microtubule bundling and recruits specific regulators of abscission[36, 57]. We mapped JADE1S expression relative to PRC1, KIF4A, and MgcRacGAP1 in cells progressing through mitosis into cytokinesis (Fig 1, C-E). As expected, in early anaphase PRC1 and KIF4 localized to the microtubule structures of the assembling central spindle: in anaphase and telophase -proteins concentrated to the midzone, and in late cytokinesis – to the stabilized tubulin fibers and midbody flanking zone (Fig 1, C, D). Interestingly, we found that in most of the cells from early to late cytokinesis, JADE1S was partially colocalized with PRC1 and KIF4 (Fig 1, C,D, columns *5-7*). In late cytokinesis JADE1S and KIF4 were localized to the same plane of the midbody flanking zone (Fig 1, D, column *8*). In late telophase the midzone JADE1S was positioned proximal to PRC1 and KIF4, but distal to AURKB (Fig 1, B-D, compare images in column *7*). Spatial-temporal expression of MgcRacGAP1 relative to JADE1S was similar to that of PRC1 (Fig 1, E). Based on plot profile analysis of late cytokinesis cells, JADE1S appears to localize precisely between the components of the two major central spindle assembly complexes and cytokinesis factors, AURKB and PRC1 (Fig 2, A-D). Our data show potential interactions with CPC and MAPs.

### 4. Central spindle targeting of JADE1S: from early midzone to late midbody

The occupancy levels of endogenous JADE1S in midzone and midbody structures of dividing cells have not been assessed. The initial analysis of asynchronously dividing cells demonstrates that endogenous JADE1S is present in the midzone of 88% of late telophase and the midbody flanking zone of 30% of late cytokinesis cells (Fig 3, A). To quantify midzone and midbody occupancy of JADE1S after mitosis exit we enriched sub-populations of mitotic and cytokinesis cells by cell cycle synchronization with mimosine (early S-phase arrest and cell cycle release), using our previously described conditions[5, 9]. Parallel samples included asynchronously dividing cells. As expected, the time– dependent increase of anaphase and telophase cell fractions started at 16 hours and reached maximum at 18 hours after cell cycle release, manifesting progression through mitosis (Fig 3 B, and not shown). Next, for the purpose of analysis, anaphase and cytokinesis cells were divided into subpopulations according to their morphology as described: anaphase A (ANA-A), anaphase B (ANA-B), early cytokinesis (early-CK), mid-cytokinesis (mid-CK), and late cytokinesis (late-CK) (see Fig 3, C for example images and description). Cells positive for JADE1S in the midzone, midbody flanking zone, and midbody were quantified within each of the sub-populations. Our results show that JADE1S association with the central spindle structures increased from 20% in anaphase A to 80.2% in anaphase B, reaching 90.3% in early cytokinesis (Fig 3, C). These data indicate that during the mitotic exit JADE1S protein is recruited to the microtubule-based structures of the forming central spindle: the midzone and the midbody flanking zone. The antibodies against JADE1 isoforms used in this study have been described and their specificity validated previously[5]. To additionally prove that antibodies used in Fig 1-4 are isoform-specific and do not cross-react we show that the recombinant JADE1L is not recognized by JADE1S antibody while the recombinant JADE1S is not detected by JADE1L antibody (Fig 3 D)[5].

### 5. Recombinant JADE1S is targeted to the inside of the midbody

We noted a consistent 50% decrease of JADE1S occupancy in midbody flanking zone of cells with particularly delayed cytokinesis (very elongated bridges). This may have resulted from either JADE1S departure from the midbody flanking zone, or, alternatively from masking inside of the dense midbody. Indeed, we were able to detect limited quantities of long cytokinesis bridges with endogenous JADE1S inside the midbody, where JADE1S nicely colocalized with KIF4A (Fig 4). It has been reported that the high density inside the midbody could mask antigens, thereby interfering with the immune-detection assays[38]. Thus, PRC1 and KIF4A became detectable inside the midbody only after addition of GFP tag to the protein[38]. Importantly, similar to PRC1, addition of GFP-tag dramatically enhanced detectability of the recombinant GFP-JADE1S protein inside the midbody in late cytokinesis cells (Fig 5, A, C). HA- and FLAG-tagged JADE1S recapitulated GFP-JADE1S, supporting localization of endogenous JADE1S to the inside of the late midbody (Fig 5, A, E).

Despite the limited quantities, we noted that, as judged by DAPI detection, all delayed cytokinesis bridges with endogenous JADE1S inside the midbody were positive for chromatin bridges, which opens an attractive possibility for JADE1S role, perhaps in unresolved chromatin detection (Fig 4, and not shown). However, considering the lower DAPI sensitivity and midbody antigen masking, the linking of JADE1S with unresolved chromatin bridges would need direct sensitive approaches.

While *per se* midbody localization of recombinant JADE1S has been documented it was not quantified and related to the ability to arrest late cytokinesis. We quantified midbody occupancy of recombinant tagged-JADE1S proteins and compared with the effects on late cytokinesis cells fraction. The effects of all three tagged JADE1S proteins on late cytokinesis cell fraction nicely correlated with their midbody occupancy, which further supported the role of midbody JADE1S in late cytokinesis arrest and abscission delay (Fig 5, B).

In sum, the dynamics of JADE1S expression during mitosis-to-cytokinesis transition shows orderly spatial interactions with microtubule-based structures of the central spindle and implicates active stepwise protein recruitment to the early midzone in anaphase, the midbody flanking zone and the midbody in mid and late cytokinesis. Analysis of JADE1S, CPC and MAP spatial-temporal expression suggest potential interactions in cytokinesis pathway.

### 6. JADE1S PHD zinc fingers are required for late cytokinesis arrest but not essential for midbody targeting

The most notable feature of JADE1 polypeptide is the presence of the two mid-molecule PHD zinc fingers. The PHD module includes the canonical PHD1 and non-canonical extended PHD2. Role of PHD2 in targeting of HAT HBO1 to chromatin and enabling acetylation of K5/12/8 lysine residues on histone H4 has been proposed[3, 7, 16]. The less studied JADE1 PHD1 might have preference to bind histone H3K36Me3 epigenetic mark, although the entire double PHD zinc finger module seems to lose this preference. Thus, the PHD module is important for JADE1S association with the bulk chromatin[6, 16].

We investigated the role of PHD zinc finger module in JADE1S cytokinesis effects. Increasing concentration of cDNA encoding the wild type or mutant JADE1S, missing the two PHD zinc fingers (JADE1Sdd) were transfected to the cells and late cytokinesis cells analyzed and quantitated. As expected, JADE1S over-expression increased late cytokinesis cell proportion in dose-dependent fashion (Fig 6, A, B). Unlike the wild type JADE1S, the mutant JADE1S_dd_ protein failed to affect scores of cytokinesis cells even when expressed at levels higher than the wild type counterpart. Similar to JADE1S, JADE1S_dd_ did not affect proportion of polynuclear cells (not shown). To examine whether the lack of PHD fingers may have affected JADE1S_dd_ midbody localization, we compared JADE1S_dd_ and JADE1S midbody occupancy in late cytokinesis cells. Interestingly, JADE1S_dd_ midbody occupancy was similar to that of the wild type HA- and FLAG-tagged JADE1S proteins (Fig 6, C). The PHD finger deletion neither affected JADE1S localization to midbody nor apparent visible morphology of cytokinesis bridge (Fig 6, D). Thus, chromatin-binding PHD zinc fingers are not essential for midbody localization but are required for JADE1S abilities to increase late cytokinesis cell proportion, suggesting the role of PHD zinc fingers in cytokinesis delay.

**Fig 6.**
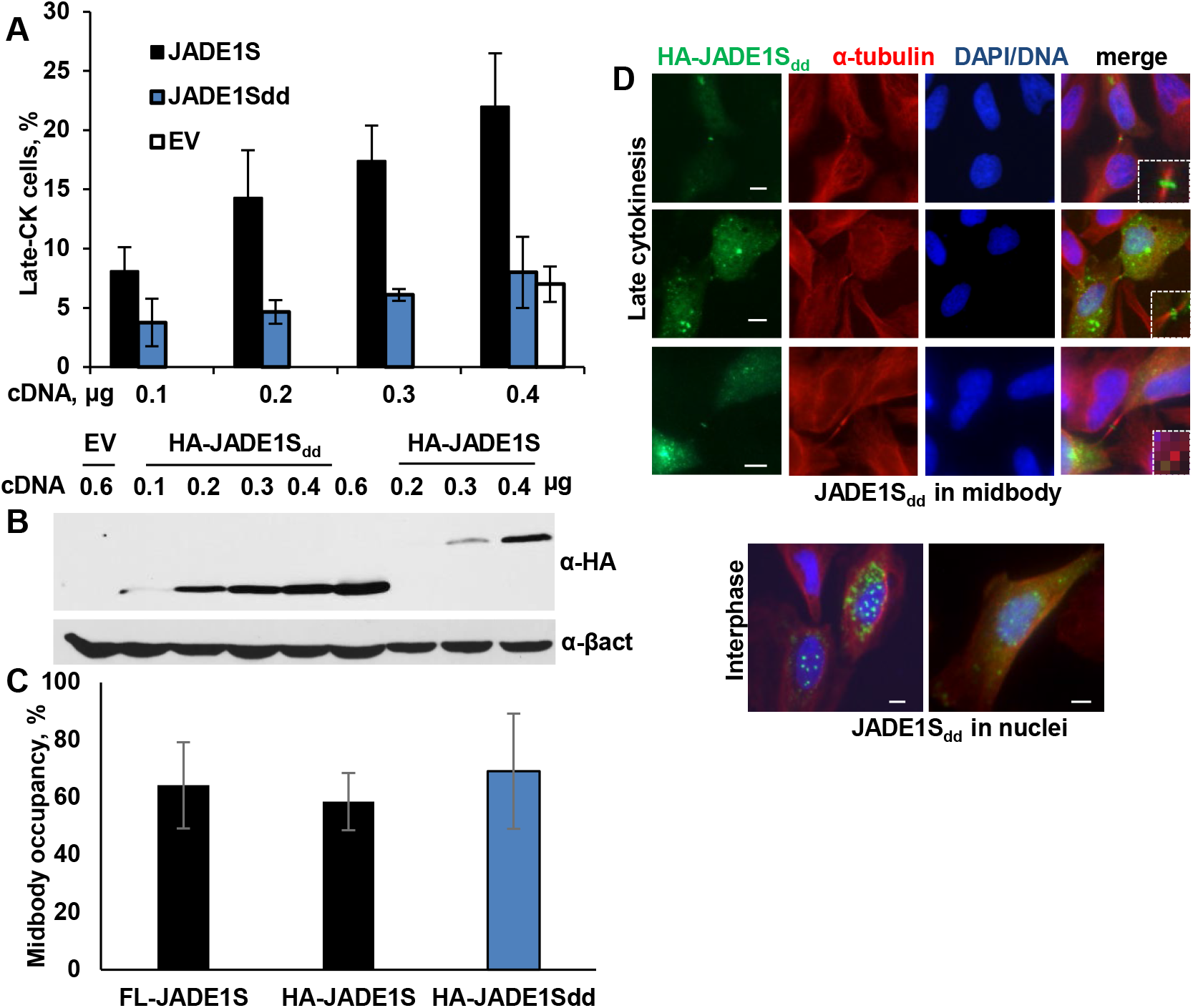
Deletion of PHD zinc finger module does not affect the midbody localization but abolishes JADE1 arrest in late cytokinesis. **(A) Effects of HA-JADE1S and HA-JADE1_dd_ on late cytokinesis cell proportion.** Dose response. Cells grown in 4-well chamber slides were transfected individually with cDNAs as indicated. Experiment was done as described in Fig 3, A,B. Values represent a percentage of the transfected late cytokinesis cells from the total number of transfected cells; three independent experiments ± s.d., >200 cells counted per transfection concentration per experiment, are included. **(B) Levels of protein expression** used in (A). HeLa cells grown in 6-well plates were transfected as indicated, proteins extracted, samples analyzed by western blots with HA-tag monoclonal antibodies and consequently β-actin. **(C) PHD fingers deletion did not affect JADE1S**_**dd**_ **midbody occupancy**. HeLa cells grown in 2-well chamber slides, were transfected with either of the cDNA: FL-JADE1S, HA-JADE1S, or HA-JADE1S_dd_ as described in (A). Proteins visualized by IF with FLAG-or HA-specific monoclonal antibody in combination with rabbit α-tubulin antibody. Total number of late cytokinesis cells expressing recombinant protein were counted, proportion of late cytokinesis cells positive for midbody recombinant protein quantified (midbody occupancy). At least 80 late cytokinesis cells total were analyzed per plasmid transfection, three independent experiments. **(D) JADE1S**_**dd**_ **is localized to the midbody in cytokinesis** (upper panels) and predominantly to the nuclei in interphase (lower panels). Representative images. Scale bar, 5 μm

### 7. HBO1 antagonizes JADE1S activities in cytokinesis

We and others previously characterized strong physical and functional interactions of nuclear JADE1S with HAT HBO1 in vitro and in life cells(7,16). According to our recent study endogenous HBO1 did not localize to late midbodies. However, neither spatial temporal expression of HBO1 in cells exiting mitosis nor effects of recombinant HBO1 on late cytokinesis cell scores has been investigated. We therefore examined potential associations of HBO1 with central spindle as well as effects on late cytokinesis cell proportion. In contrast to JADE1S, endogenous as well as recombinant HBO1 did not localize to the central spindle structures, including midzone and midbody, in cells exiting mitosis and undergoing anaphase and cytokinesis (Fig 7). Unlike JADE1S, the recombinant HBO1 protein did not increase late cytokinesis cell fraction (Fig 8, A). In fact, even the lowest quantity of transfected HBO1 protein resulted in an opposite effect and decreased the basal levels of late cytokinesis cells (Fig 8, A). Hence, we hypothesized that HBO1 might interfere with JADE1S activities on cytokinesis arrest. Indeed, according to co-transfection experiments, the expression of HBO1 completely prevented JADE1S-mediated increase of late cytokinesis cell proportion. To determine whether enzymatic activity of HBO1 plays a role in attenuation of this JADE1S-mediated cytokinesis effect, we took advantage of the previously characterized mutant of HBO1 which physically interacts with JADE1S but lacks the enzymatic activity. Similar to the wild type counterpart, the acetylation dead mutant of HBO1 decreased basal levels of late cytokinesis cells, and upon co-transfection, prevented JADE1S-mediated increase of late cytokinesis cell fraction. Thus, the negative effect of HBO1 did not depend on enzyme activity (Fig 8, B).

**Fig 7.**
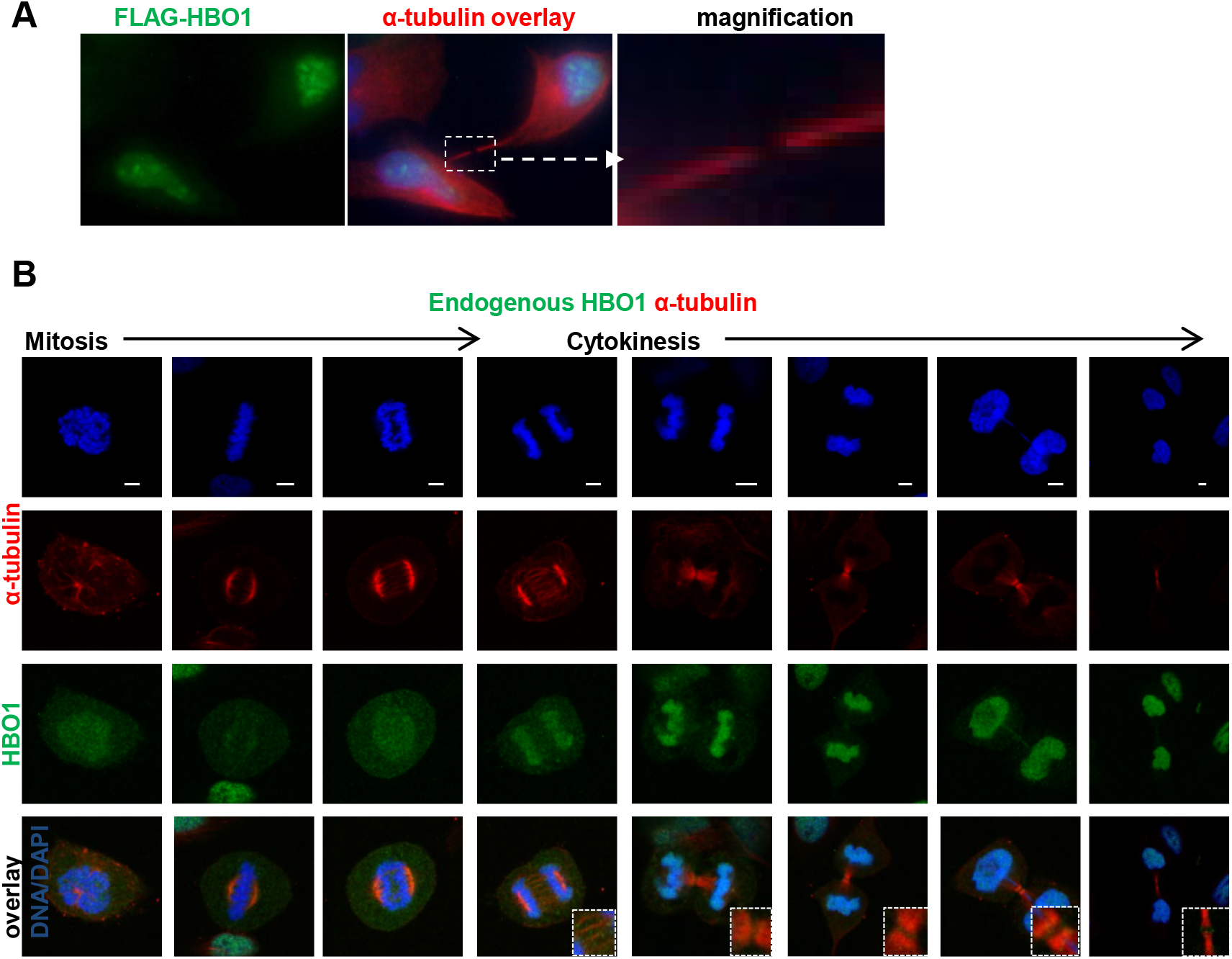
FLAG-HBO1 and endogenous HBO1 are not expressed in midbody and midzone. (**A**) FLAG-HBO1 and (**B**) endogenous HBO1 in HeLa cells. Immunofluorescence was performed as described in Fig 1 and in Methods, except FLAG monoclonal and HBO1 polyclonal affinity purified antibody were used. DAPI counterstain. Scale bar, 5 μm

**Fig 8.**
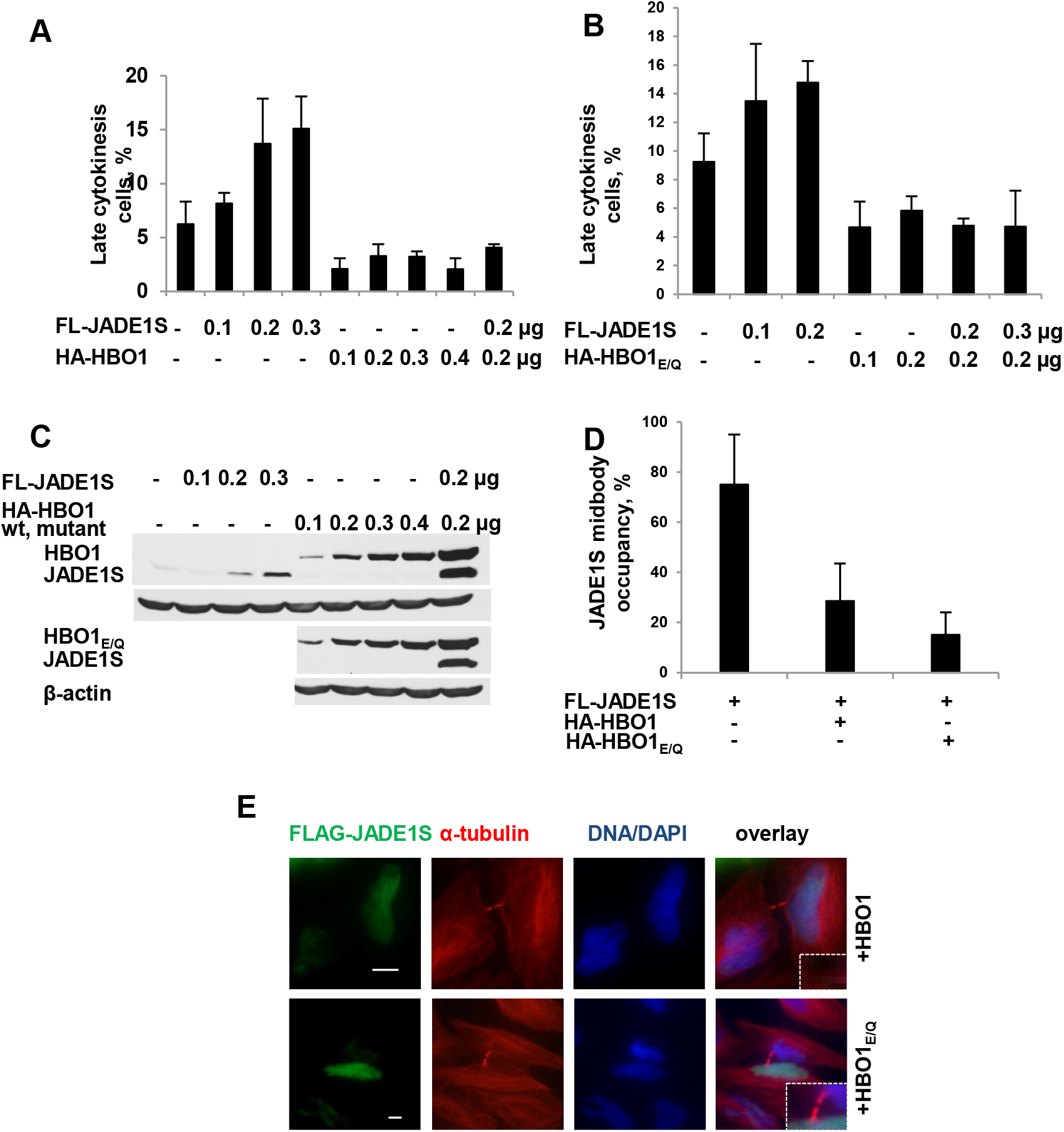
Recombinant HBO1 decreases late cytokinesis cells proportion, abolishes recombinant JADE1S-mediated late cytokinesis arrest and prevents JADE1S midbody localization. **(A, B)** HeLa cells grown in two-well chamber slides were transfected individually or in combination with cDNA for FLAG-JADE1S, HA-HBO1 and HA-HBO1_E/Q_ and late cytokinesis cells proportion quantitated as described in Figs 3 and 4. Empty vector plasmid was included in control (none) and to fill in cDNA. For FLAG-JADE1S at least 80 late cytokinesis cells were analyzed per experimental point, for wild type or mutant HBO1 at least 40 cells were analyzed. Experiments were repeated twice. **(C)** Western blot shows levels of expression of the effectors used in (A,B). Experiment was done as described in Fig 4. HeLa cell grown in 6-well plates were transfected as indicated, proteins extracted and analyzed for transgene expression. Same nitrocellulose membrane was sequentially probed with FLAG-, HA-tag, and β-actin monoclonal antibodies. **(D, E)** Wild type HBO1 and mutant HBO1_E/Q_ both decrease the rates of JADE1S midbody occupancy. Cells grown on two-well chamber slides were transfected with 0.2μg of plasmids as indicated, FLAG-JADE1S and β-actin visualized by immunofluorescence staining. Single cell co-expression and co-localization of recombinant proteins (wild type and mutant HA-HBO1 and FLAG-JADE1S) was verified in parallel samples, the co-expression detected in >90% of transfected cells by using a combination of HA- and FLAG-antibodies (not shown and in previous study(9)). Representative images (E). Scale bar, 10μm

Next, we examined whether by interacting with JADE1S, HBO1 might interfere with JADE1S midbody localization. Indeed, expression of recombinant HBO1 effectively decreased JADE1S midbody occupancy (Fig 8, D). Interestingly, similarly to the wild type HBO1, the enzymatically inactive mutant HBO1_E/Q_ also decreased fraction of late cytokinesis cells with the midbody JADE1S. We hypothesize that HBO1 interferes with JADE1S cytokinesis arrest by affecting JADE1S compartmentalization and preventing midbody targeting.

## Discussion

The current study revealed several insights about JADE1S expression and function during cytokinesis progression: requirement of JADE1S PHD fingers, antagonistic relationship with HBO1 partner, and early central spindle targeting of JADE1S. Topological analysis of JADE1S expression showed that JADE1S is directed to the microtubule-based structures of the central spindle beginning as early as in anaphase. In cells exiting mitosis JADE1S concentrated to the early midzone, followed by late midzone, the midbody flanking zone, and mature midbody in late cytokinesis. Quantitative analyses of JADE1S occupancy rates with these central spindle structures indicated step wise recruitment. Moreover, analysis of plot profiles of JADE1S and components of CPC, MAPs, and centralspindlin revealed transient colocalization in telophase and mapped central spindle location of JADE1S between AURKB (CPC) and PRC1-KIF4A (MAP). According to our recent study recombinant JADE1S-mediated late cytokinesis arrest is released by AURKB inhibitor, while JADE1S depletion increased polynucleation and aneuploidy. Results of topological analysis of JADE1S expression in cells transitioning from mitosis to early and late cytokinesis support the role of JADE1S in cytokinesis regulation pathway, suggests recruitment and interaction with established cytokinesis factors [5, 9].

According to our quantitation analysis, a large proportion of mid cytokinesis cells bear endogenous JADE1S in the midbody flanking zone. We also detected endogenous JADE1S protein inside the midbody (Flemming body) in cells with particularly delayed cytokinesis and long bridges. In those cells JADE1S clearly co-localized with Flemming body MAP protein, KIF4A. The lower percentage of Flemming body JADE1S was likely due to the antigen masking within the highly dense structure. Interestingly, unlike the endogenous protein, the recombinant GFP-JADE1S was detected inside the midbody in the vast majority of late cytokinesis cells even when expressed at the low levels. This phenomena has been described for other proteins, including alpha-tubulin, centralspindlin complex protein MKLP1, and microtubule associated proteins PRC1 and KIF4[38]. The GFP-tagged but not endogenous species of these proteins revealed their localization to the dense center of the midbody. Our topologic and quantitative analyses strongly suggest that during mitosis exit and cytokinesis, JADE1S interacts with microtubule-based structures of the central spindle in order to participate in late cytokinesis control.

The transient co-localization of JADE1S with AURKB in telophase is intriguing. One of the key kinase of late mitosis and cytokinesis, AURKB is required for the cytokinesis progression[42, 58]. Importantly, according to the recent discovery, AURKB is also required for the final abscission checkpoint[47-49]. The checkpoint could be activated by fibers of unresolved chromatin caught within the cytokinesis bridge (also called lagging or dragging chromatin, chromatin bridges). We recently communicated that increase of late cytokinesis cell proportion by recombinant JADE1S is reversed by the addition of AURKB pharmacological inhibitor. Earlier we reported that JADE1S protein is transiently phosphorylated in early mitosis, and identified six phosphorylated residues by Mass spectrometry. Together with previous studies, the co-localization results of this study further support that JADE1S might be a target of AURKB to function in final abscission delay[5, 9]. Given PHD zinc finger-dependent chromatin interactions of JADE1, it’s tempting to speculate that AURKB facilitates JADE1S recruitment to the central spindle and the midbody to bind or perhaps recognize the unresolved chromatin as part of delay. In fact, this would agree with the result showing that PHD zinc finger deletion yielded a mutant incapable of increasing late cytokinesis cell proportion even when expressed at high levels. Note, that the removal of the PHD zinc fingers did not affect JADE1S midbody localization, showing that JADE1S effects on cytokinesis delay but not the midbody targeting required PHD module.

Little is known about the cell cycle effects of interactions between JADE1S and HBO1. We show that the addition of low levels of HBO1 protein to the dividing cells prevented increase of late cytokinesis cells proportion caused by recombinant JADE1S. The effect did not depend on the HAT activity as enzymatically inactive HBO1_E/Q_ fully mimicked the effects of the wild type HAT. Because 1) HBO1 co-transfection greatly reduced JADE1S midbody occupancy while increasing the total JADE1S abundance, and 2) HBO1 alone decreased basal levels of late cytokinesis fraction, it is possible that the antagonistic effect of HBO1 is due to withdrawal of JADE1S from the central spindle, specifically from the midbody. More studies will clarify this possibility.

The results of this study shed light on the spatial-temporal regulation of JADE1S during cytokinesis progression and suggested interactions with central spindle factors, including key CPC component, AURKB. We identified that JADE1S PHD zinc finger module is required for cytokinesis arrest function, but is dispensable for JADE1S midbody targeting. Further, our co-transfection experiments identified relationship with JADE1S partner, HAT HBO1. The negative effects of recombinant HBO1 on JADE1S-mediated late cytokinesis arrest and its midbody localization show functional interactions and suggest an antagonistic role. The results of this study support role JADE1S in cytokinesis delay.

## Acknowledgement

Authors are grateful to Rytis Prekeris for reading the manuscript and providing valuable feedback. This work was supported by National Institutes of Health Grant RO1 DK087910.

